# Direct Injection NanoHILIC/MS/MS Proteomics from Reversed-Phase StageTip Eluate

**DOI:** 10.64898/2026.05.27.728107

**Authors:** Koshin Akamatsu, Eisuke Kanao, Yasushi Ishihama

**Affiliations:** Graduate School of Pharmaceutical Sciences, Kyoto University, Kyoto 606–8501, Japan

**Author notes:** Correspondence and requests for materials should be addressed to E.K. and Y.I. Eisuke Kanao, Yasushi Ishihama.

**Keywords:** nanoHILIC/MS/MS, bottom-up proteomics, peptide solubility, direct injection

## Abstract

NanoHILIC/MS/MS provides high sensitivity for low-input peptide analysis, yet its use in bottom-up proteomics has been constrained by a persistent solvent mismatch: tryptic peptides exhibit poor solubility in the ≥95% acetonitrile (ACN) required for nanoHILIC injection. Our previously reported two-step solubilization method [*Anal* Chem 2025, 97 (19), 10227–10235] alleviated this issue but required large dilution volumes, limiting the amount of sample that could be injected. Here, we introduce DiReCT (Dissolution from Reverse-Phase Chromatography Tips), a StageTip-based workflow that integrates peptide solubilization, desalting, and nanoHILIC-compatible elution into a single operation. During elution from RP-StageTips, residual water on the stationary phase is rapidly displaced by a small volume of high-ACN solvent, generating a transient mid-ACN environment that maximizes peptide solubility without drying. This mechanism enables high-recovery peptide concentration and allows direct injection of the entire eluate onto nanoHILIC/MS/MS. Using ∼0.25 ng of HeLa digest, DiReCT/nanoHILIC/MS/MS identified 1177 peptides and 410 proteins, representing 8.9- and 6.7-fold increases over nanoRPLC/MS/MS, respectively.

## INTRODUCTION

Bottom-up proteomics based on nanoscale liquid chromatography–tandem mass spectrometry (nanoLC/MS/MS) has become an essential analytical platform for comprehensive characterization of cellular proteins and their functional states.^1,2^ Recent advances in MS instrumentation—particularly ultrafast acquisition exceeding 200 Hz and markedly improved sensitivity—have accelerated applications such as high-throughput single-cell proteomics^3,4,5,6^ and clinical spatial proteomics.^7^ Despite these developments, liquid chromatography (LC) remains a critical determinant of identification performance and throughput.^8–11^ Improvements in LC have been achieved through the development of monolithic^12,13^ and pillar-array columns,^14^ optimization of mobile-phase composition,^15–17^ and integration with automated sample-preparation systems.^18,19^ However, most advances have remained within the framework of nanoscale reversed-phase LC (nanoRPLC).^8^ Fully leveraging the speed and sensitivity of state-of-the-art MS requires reconsideration of the separation mode itself.

Nanoscale hydrophilic interaction liquid chromatography (nanoHILIC)/MS/MS has emerged as a promising alternative to nanoRPLC/MS/MS. In contrast to RPLC, HILIC employs polar stationary phases and mobile phases containing high proportions of acetonitrile (ACN), providing separation selectivity largely orthogonal to RPLC.^20–22^ The high ACN content reduces solvent viscosity and column pressure, enabling faster separations,^23–25^ while also enhancing droplet desolvation and ionization efficiency in electrospray ionization (ESI), thereby improving MS sensitivity.^22,26^ We previously demonstrated that nanoHILIC/MS/MS using silica monolithic columns achieves higher sensitivity and a wider dynamic range than nanoRPLC/MS/MS.^27^

A major practical challenge in nanoHILIC, however, is its pronounced sensitivity to the composition and volume of the injection solvent. Because tryptic peptides exhibit poor solubility in ≥95% ACN—the solvent required for stable HILIC retention—samples must typically be dried and redissolved in a minimal volume of the initial mobile phase. This solvent mismatch between peptide solubilization and HILIC injection severely limits sensitivity. To address this issue, we recently developed a two-step solubilization method in which peptides are first dissolved in 25% ACN and then diluted into 95% ACN.^28^ This approach improved peptide recovery and significantly enhanced nanoHILIC/MS/MS performance relative to nanoRPLC/MS/MS. However, the large dilution factor resulted in a relatively high final sample volume, restricting the amount of material that could be injected and preventing seamless integration of post-digestion desalting with HILIC-compatible sample introduction.

Injection strategies such as at-column dilution (ACD) and performance-optimizing injection sequence (POISe) have been proposed for larger-bore HILIC systems to mitigate solvent mismatch by diluting the sample plug immediately before or during injection.^29–31^ In nanoHILIC, however, the internal column volume is only ∼1.5 μL, making analyte retention highly sensitive to even slight decreases in ACN content.^28^ As a result, ACD and POISe are difficult to implement without substantial sample dilution or extremely small injection volumes, both of which compromise low-input analysis. We therefore reasoned that resolving solvent mismatch upstream—during sample preparation rather than at injection—would be more effective for nanoHILIC/MS/MS.

Here, we introduce DiReCT (Dissolution from Reverse-Phase Chromatography Tips), a sample-preparation workflow based on stop-and-go extraction tips (StageTips).^32,33^ In DiReCT, peptides are acidified after trypsin digestion, loaded onto an RP-StageTip, and concentrated. During elution, residual water on the stationary phase is rapidly displaced by a small volume of high-ACN solvent, generating a transient mid-ACN environment that maximizes peptide solubility without drying. This design enables high-recovery peptide concentration, integrates desalting and solubilization into a single operation, and yields eluates directly compatible with nanoHILIC injection. Together, these features overcome the long-standing solvent mismatch in nanoHILIC sample preparation and expand the practical applicability of nanoHILIC/MS/MS to low-input proteomics.

## MATERIALS AND METHODS

### Materials

UltraPure Tris buffer was obtained from Thermo Fisher Scientific (Waltham, MA). Sequencing-grade modified trypsin was purchased from Promega (Madison, WI). Water was purified using a Milli-Q system (Millipore, Bedford, MA). Empore SDB-XC disks were purchased from GL Sciences (Tokyo, Japan). Blunt-end 16- and 20-gauge syringe needles were obtained from Hamilton (Reno, NV). Protease and phosphatase inhibitors were purchased from Sigma-Aldrich (St. Louis, MO). Unless otherwise noted, all chemicals were from Fujifilm Wako (Osaka, Japan). Pipette tips and microtubes (Eppendorf, Hamburg, Germany) were used for sample handling, and Axygen™ 96-well PCR plates (Corning, NY) were used for sample storage. A Se-08 vortex mixer (TAITEC, Saitama, Japan) and a UCT-1331 ultrasonic bath (TOCHO, Tokyo, Japan) were used during sample preparation.

### Cell culture

HeLa S3 cells (JCRB Cell Bank, Osaka, Japan) were cultured in Dulbecco’s modified Eagle’s medium supplemented with 10% fetal bovine serum in 10-cm dishes until 80% confluency. Cells were washed twice with ice-cold PBS, collected using a cell scraper, and pelleted by centrifugation.

### Protein extraction and protease digestion

Proteins were extracted using a phase-transfer surfactant protocol as described previously.^34^ Cell pellets were suspended in lysis buffer containing 100 mM Tris-HCl (pH 9), protease inhibitors, phosphatase inhibitor cocktails 2 and 3, 12 mM sodium deoxycholate, and 12 mM sodium lauroyl sarcosinate. The suspension was heated at 95 °C for 5 min and sonicated for 20 min. Protein concentration was determined using a BCA assay.

Proteins were reduced with 10 mM dithiothreitol for 30 min and alkylated with 50 mM iodoacetamide for 30 min in the dark. Samples were diluted 5-fold with 50 mM ammonium bicarbonate and digested with Lys-C (1:100, w/w) for 3 h at room temperature, followed by overnight trypsin digestion (1:100, w/w) at 37 °C. BSA was processed identically. After digestion, 1 mL ethyl acetate was added to 1 mL digest, and the mixture was acidified to 0.5% TFA. After vortexing (2 min) and centrifugation (15,800 × g, 2 min), the aqueous phase was collected, pooled, aliquoted, and stored at −80 °C.

### DiReCT method

Peptide samples were prepared in 0.5% TFA in water (buffer A). RP-StageTips were prepared as described previously.^32,33^ Empore SDB-XC disks (0.5-mm thickness) were punched using 16- or 20-gauge needles to obtain 1.2- or 0.6-mm disks, respectively, and inserted into 200-µL or 10-µL pipette tips. All StageTip operations were performed at 4 °C by centrifugation to minimize evaporation.

StageTips were conditioned with 20 µL of buffer B (95% or 99.9% ACN with 0.1% FA) followed by 20 µL of buffer A. Peptide samples in buffer A were loaded, washed with 20 µL of buffer A, and eluted with buffer B into a 96-well plate. Eluates were injected directly onto the nanoHILIC/MS/MS system.

### Two-step solubilization method

As a control, the two-step solubilization method was performed as described previously.^28^ Pre-desalted peptides (5 µg) in 0.5% TFA were processed using a StageTip, collected, and dried in a SpeedVac. Dried peptides were dissolved in 4 µL of 25% ACN with 0.1% FA and diluted with 96 µL of 97.9% ACN with 0.1% FA to yield 95% ACN.

### Dried-sample comparison between DiReCT and the two-step solubilization method

To eliminate drying-related effects, 5-µg aliquots of peptides were dispensed into 96-well plates and dried. For the two-step method, dried peptides were dissolved and diluted as above. For DiReCT, a StageTip with one 1.2-mm SDB-XC disk was equilibrated with buffer B and buffer A. Dried peptides were dissolved in 4 µL of 25% ACN with 0.1% FA, mixed with 20 µL of buffer A in the StageTip body, and loaded by brief centrifugation without drying the disk. The well was rinsed with an additional 4 µL of 25% ACN and 20 µL of buffer A, and the rinse was loaded onto the same StageTip. After washing with 20 µL of buffer A, peptides were eluted with 100 µL of 95% ACN with 0.1% FA. A 5 µL aliquot was analyzed by nanoHILIC/MS/MS on a Q-Exactive mass spectrometer (Thermo Fisher Scientific). Detailed Q-Exactive acquisition parameters are available in the Supporting Information.

### Estimation of the residual water amount in a StageTip

Experiments were performed at 4 °C to prevent evaporation. Ten RP-StageTips containing one 1.2-mm SDB-XC disk were weighed dry, conditioned with 20 µL ACN followed by 20 µL water, centrifuged, and weighed again. Residual water per StageTip was calculated from the mass difference.

### NanoHILIC/MS/MS

NanoHILIC/MS/MS was performed on a timsTOF HT mass spectrometer (Bruker Daltonics, MA) coupled to an UltiMate 3000 nanoLC system (Thermo Fisher Scientific) and a PAL HTC-xt autosampler (CTC Analytics, Zwingen, Switzerland). Peptides were separated on a self-pulled 200-mm × 100-µm i.d. needle column packed with ZIC-HILIC particles (3.5 µm, 100 Å, Merck, Darmstadt, Germany) at 1000 nL/min and 25 °C. Mobile phase A was 0.1% FA in ACN; mobile phase B was 0.1% FA in water. The gradient was: 0–5 min, 5–20% B; 5– 25 min, 20–35% B; 25–40 min, 35–60% B; 40–45 min, 60–70% B; 45–45.1 min, 70–5% B; 45.1–60 min, 5% B. Injection volume was 5 µL. Columns were prepared according to a previously described procedure.^35^ Briefly, fused-silica capillaries were pulled using a P-2000 laser puller (Sutter Instruments, Novato, CA). A slurry of ZIC-HILIC particles in methanol was packed into the capillaries under 10 MPa N_2_ pressure.

MS data were acquired in ddaPASEF mode.^36^ One TIMS-MS scan was followed by 10 PASEF MS/MS scans. The *m/z* range was 100–1700; ion mobility range was 1/K_0_ = 0.6–1.5 V·s cm^−2^. Singly charged ions were excluded. Precursors above 2500 a.u. were selected and dynamically excluded for 0.4 min. Quadrupole isolation widths and collision energies were set as previously described.^28^ The electrospray source was operated at 1400 V with 3.0 L/min dry gas at 180 °C.

### NanoRPLC/MS/MS

NanoRPLC/MS/MS was performed on the same LC–MS platform. Peptides were separated on a 200-mm × 100-µm i.d. self-pulled needle column packed with ReproSil-Pur 120 C18-AQ particles (3 µm, 120 Å). Mobile phase A was 0.1% FA in water; mobile phase B was 80% ACN with 0.1% FA. The gradient was: 0–40 min, 5– 40% B; 40–42 min, 40–99% B; 42–45 min, 99% B; 45–45.1 min, 99–5% B; 45.1–60 min, 5% B. Injection volume was 5 µL; flow rate was 1000 nL/min.

### Data analysis

HeLa tryptic peptides were analyzed using FragPipe v23.1 with MSFragger v4.3 and IonQuant v1.11.11 against the UniProtKB/Swiss-Prot human database (November 2024, 20,429 entries). Mass tolerances were 20 ppm for precursors and fragments. Trypsin/P was specified with up to two missed cleavages. Carbamidomethylation of cysteine was fixed; methionine oxidation and protein N-terminal acetylation were variable. MSBooster rescoring used spectral and ion-mobility predictions. Results were filtered at 1% FDR at the PSM and protein levels. BSA peptides were quantified using Skyline-daily v23.1.1.459.^37^

## RESULTS AND DISCUSSION

### Concept and Mechanism of the DiReCT Workflow

Figure 1 summarizes the conceptual basis of the DiReCT workflow for nanoHILIC/MS/MS. After tryptic digestion, peptides are present in an acidic aqueous solution and are retained on an RP-StageTip during loading and washing. Elution with a small volume of high-ACN solvent initiates solvent exchange, enabling direct injection onto a HILIC column. As the ACN-rich eluent displaces the residual aqueous solution within the stationary phase, a transient ACN gradient is formed. Peptides experience this gradient before being recovered in a nanoHILIC-compatible small-volume eluate.

**Figure 1.**
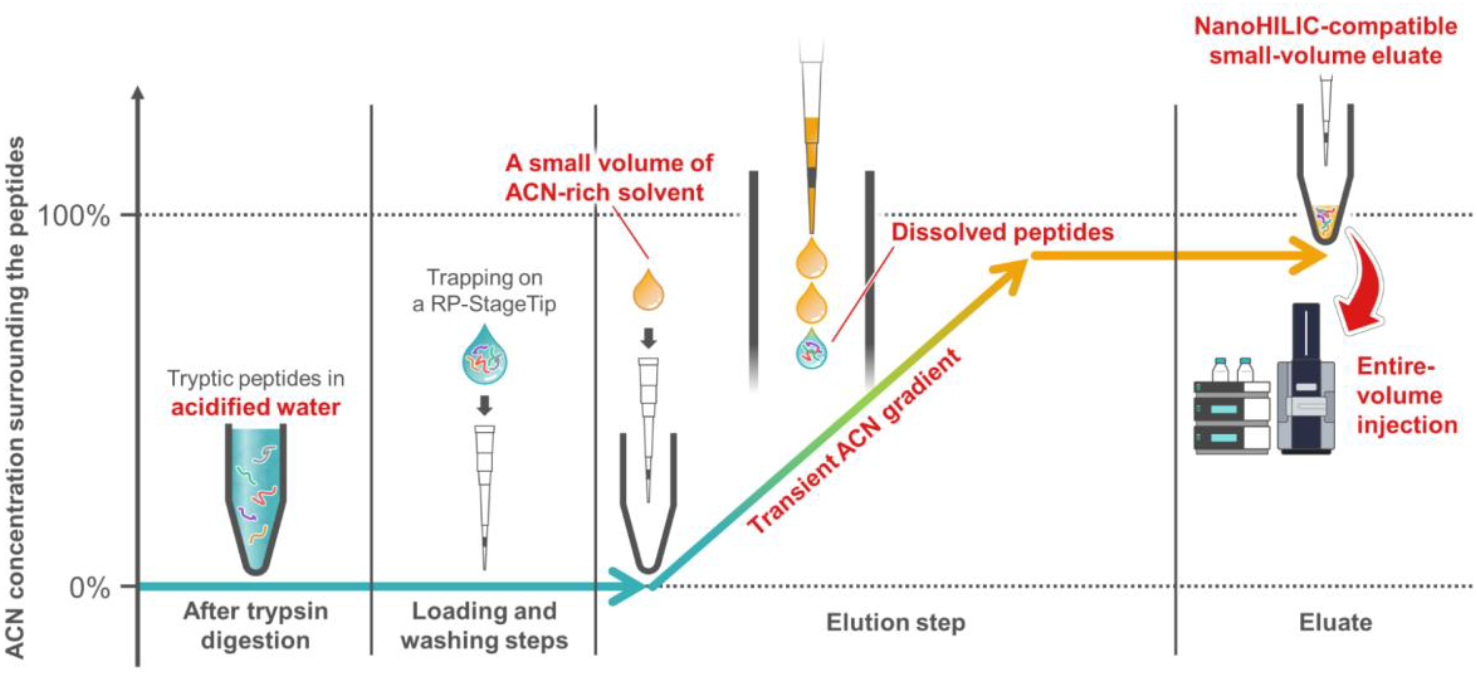
Conceptual overview of the DiReCT workflow. Peptides retained on an RP-StageTip under acidic aqueous conditions are directly eluted with a small volume of high-ACN solvent. As the ACN-rich eluent displaces residual aqueous solution within the stationary phase, peptides experience a transient ACN-composition gradient before being recovered in a nanoHILIC-compatible small-volume eluate. This process eliminates post-desalting dry-down and reconstitution and enables complete sample transfer for nanoHILIC/MS/MS.

Our previous ACN–solubility profiling of tryptic peptides showed that peptide solubility is maximized across a broad range centered around 25% ACN.^28^ The key feature of DiReCT is that it does not simply replace aqueous solvent with 95–99.9% ACN; rather, it sequentially transitions peptides through three solvent environments— post-digestion aqueous, optimal solubilization (mid-ACN), and HILIC injection (high-ACN)—within a single StageTip. This design eliminates drying and reconstitution steps and enables complete sample transfer.

### Comparison with the Two-Step Solubilization Method

#### A. Dilution-matched comparison

To compare DiReCT with the two-step solubilization method under conservative conditions, DiReCT was intentionally performed using a large elution volume (100 µL), matching the final volume of the two-step workflow. HeLa peptides (5 µg) in 0.5% TFA were prepared at 50 ng/µL, and 5 µL aliquots (250 ng) were analyzed by nanoHILIC/MS/MS. Both methods yielded comparable numbers of identified peptides with similar reproducibility (*p* = 0.056, Figure 2A). Among commonly quantified peptides, 72.9% showed higher intensities with DiReCT, with a mean fold-change of 1.22 (Figure 2B). Thus, even when DiReCT’s low-volume advantage is intentionally suppressed, StageTip-based elution provides equal or improved peptide recovery under nanoHILIC-compatible conditions.

**Figure 2.**
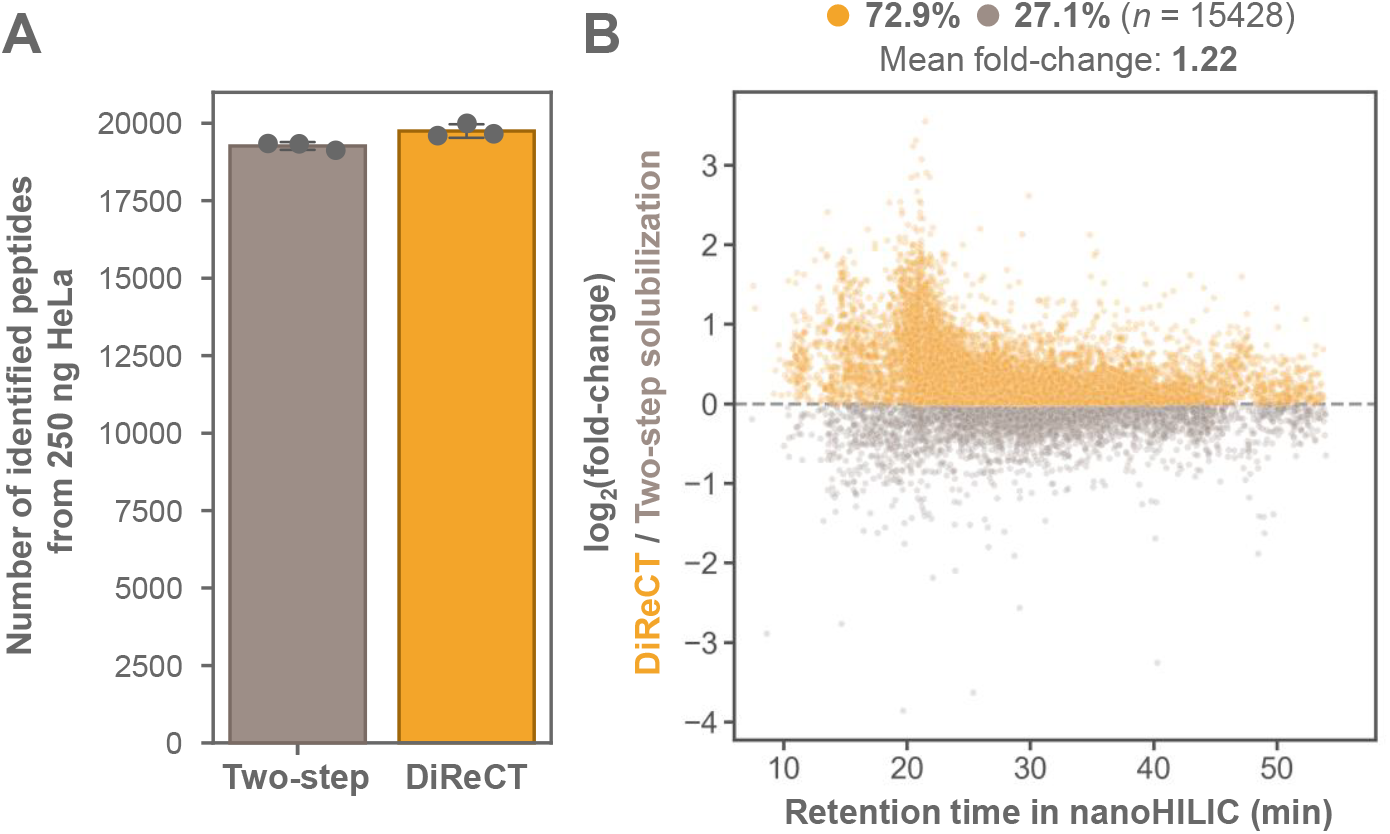
Comparison of DiReCT and the two-step solubilization method under dilution-matched (100 µL) conditions. (A) Numbers of peptides identified from 250 ng of HeLa peptides prepared by DiReCT or the two-step solubilization method. (B) Log_2_(fold-change) in intensities of commonly quantified peptides plotted against nanoHILIC retention time. Yellow and gray dots indicate peptides with higher intensities in DiReCT and the two-step method, respectively. Sample preparation was performed in triplicate.

#### B. Dried-Sample Comparison

We next evaluated whether DiReCT performs equivalently when peptides have undergone a dry-down step (Figure 3A). Dried peptides were reconstituted in 25% ACN for both workflows to match the initial condition of the two-step method. In DiReCT, the reconstituted sample was diluted with buffer A inside the StageTip, loaded, washed, and eluted. Both methods identified similar numbers of peptides from 250 ng of HeLa digest (*p* = 0.15; Figure 3B). Peptide intensities were nearly balanced, with 51.5% showing higher intensities with DiReCT and a mean fold-change of 1.04 (Figure 3C). These results indicate that the two solubilization strategies—(i) dissolving in 25% ACN followed by dilution, and (ii) eluting peptides from an RP disk via a transient ACN gradient—are mechanistically comparable in terms of recovery.

**Figure 3.**
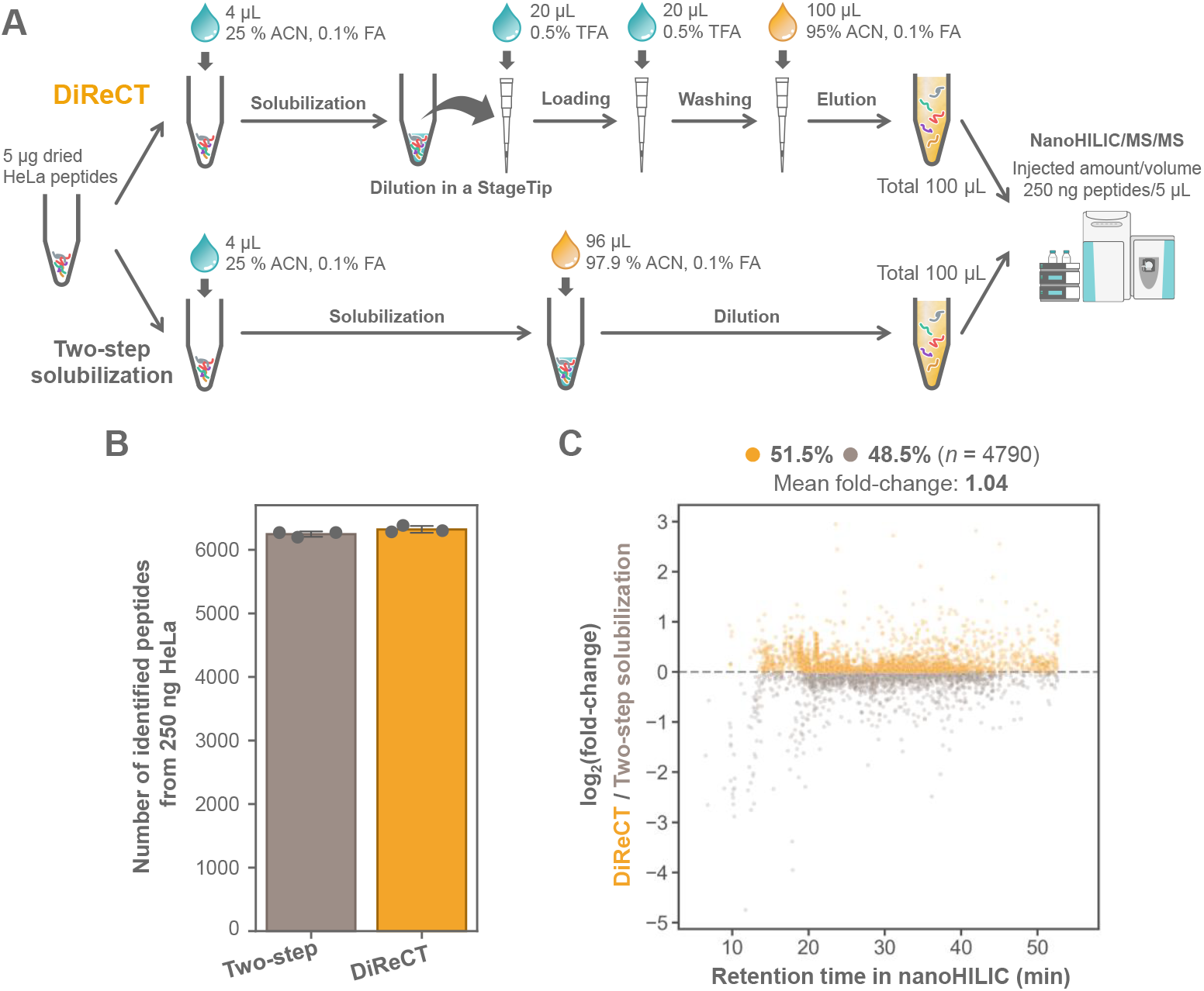
Dried-sample comparison between DiReCT and the two-step solubilization method. (A)Experimental workflow. Dried HeLa peptide aliquots (5 µg) were processed by either method under matched dissolution and final-volume conditions. (B) Numbers of peptides identified from 250 ng of peptides. (C) Log_2_(fold-change) in intensities of commonly quantified peptides plotted against nanoHILIC retention time. Yellow and gray dots indicate peptides with higher intensities in DiReCT and the two-step method, respectively. Data were acquired on a Q-Exactive platform. Sample preparation was performed in triplicate.

Together with the results in Figure 2, these findings support a model in which peptides concentrated on the RP disk under aqueous conditions can be efficiently recovered through the transient ACN gradient generated during DiReCT elution. Importantly, previously reported RP-based solvent-exchange systems for HILIC injection were developed for larger-bore LC formats and under conditions where neither sample solubility in high-ACN solvents nor complete transfer of the entire sample volume was a critical requirement.^**38**, **39**^ As such, these approaches address a fundamentally different problem and do not target the low-volume, full-injection, peptide-solubility constraints that DiReCT is designed to overcome.

### Downscaling DiReCT for Entire-Volume Injection at the 5 µL Scale

We next examined whether DiReCT could be downscaled for entire-volume injection without compromising nanoHILIC performance. Peptide concentration was fixed at 50 ng/µL, while elution volumes were reduced from 100 to 5 µL (Figure 4A). As elution volume decreased, peptide identifications declined, with the largest drop at 5 µL. Chromatographic inspection revealed reduced signal intensity, peak broadening, and breakthrough in the 5 µL format (Figure 4B, C), consistent with insufficient ACN content in the injected solvent. Gravimetric measurements showed that a 1.2-mm RP disk retained 0.44 µL of residual aqueous solvent, reducing the effective ACN concentration of a 5 µL eluent from 95% to 87.3% (Figure S1). This dilution is negligible at 100 µL but critical at 5 µL. To minimize the impact of residual aqueous solvent, we introduced two modifications: (i) increasing the ACN concentration of the eluent from 95% to 99.9%, and (ii) reducing the RP disk diameter from 1.2 mm to 0.6 mm, thereby decreasing the retained solvent volume by approximately fourfold. Together, these adjustments increased the estimated ACN concentration of the final eluate to 97.7% (Figure S1B). Under these optimized conditions, entire-volume injection of 5 µL eluates yielded peptide identifications comparable to the 100 µL format (Figure 5B, Figure S2). Chromatographic performance improved markedly, with sharper peaks and reduced breakthrough (Figure 5C, D). Among commonly quantified peptides, 56.4% showed higher intensities under optimized conditions, with a mean fold-change of 2.02 (Figure 5E).

**Figure 4.**
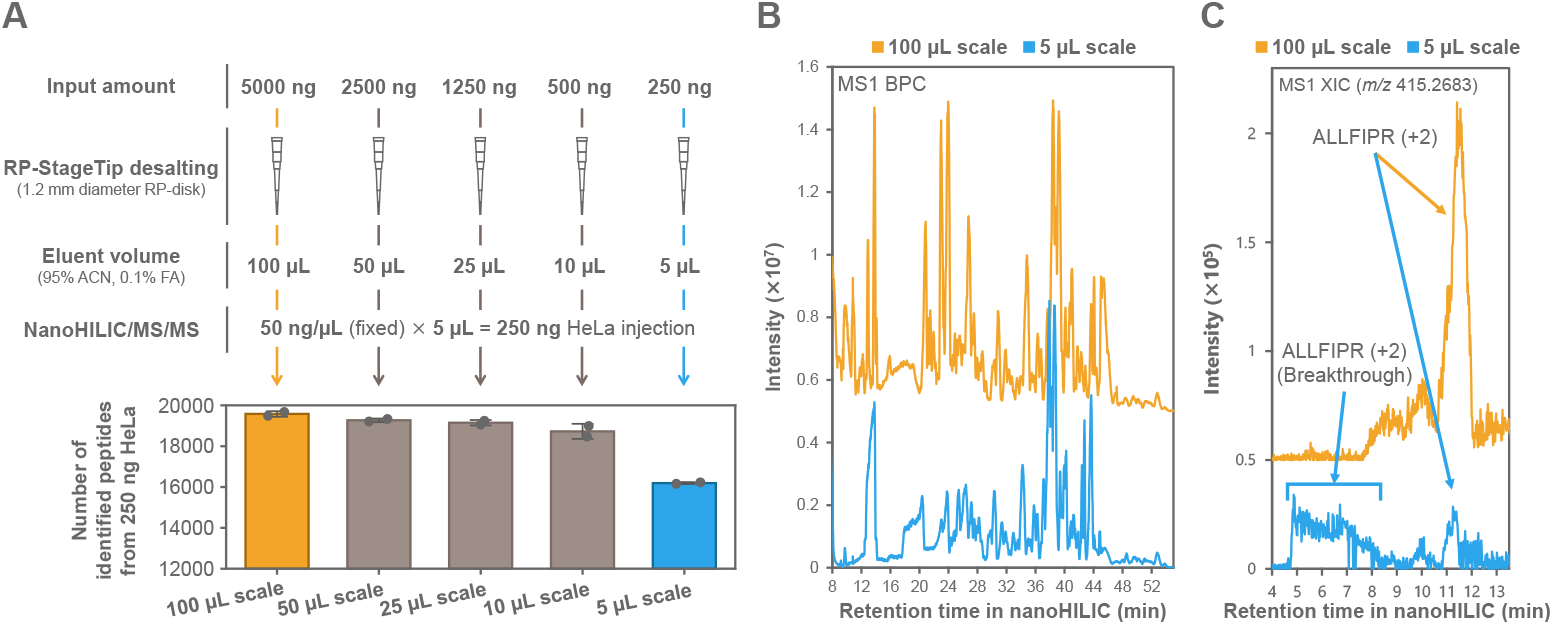
Effect of elution volume on nanoHILIC/MS/MS performance in DiReCT. (A) Overview of the experiment and numbers of identified peptides obtained from eluates ranging from 5 to 100 µL. (B) MS1 base-peak chromatograms for the 100- and 5-µL formats. (C) Extracted ion chromatograms (m/z 415.2683, peptide ALLFIPR^2+^). HeLa peptides (250–5000 ng) were desalted on RP-StageTips packed with a 1.2-mm RP disk and eluted with 5–100 µL of 95% ACN containing 0.1% FA, maintaining a final peptide concentration of 50 ng/µL. A 5-µL aliquot (250 ng) was analyzed by nanoHILIC/MS/MS. Sample preparation was performed in duplicate.

**Figure 5.**
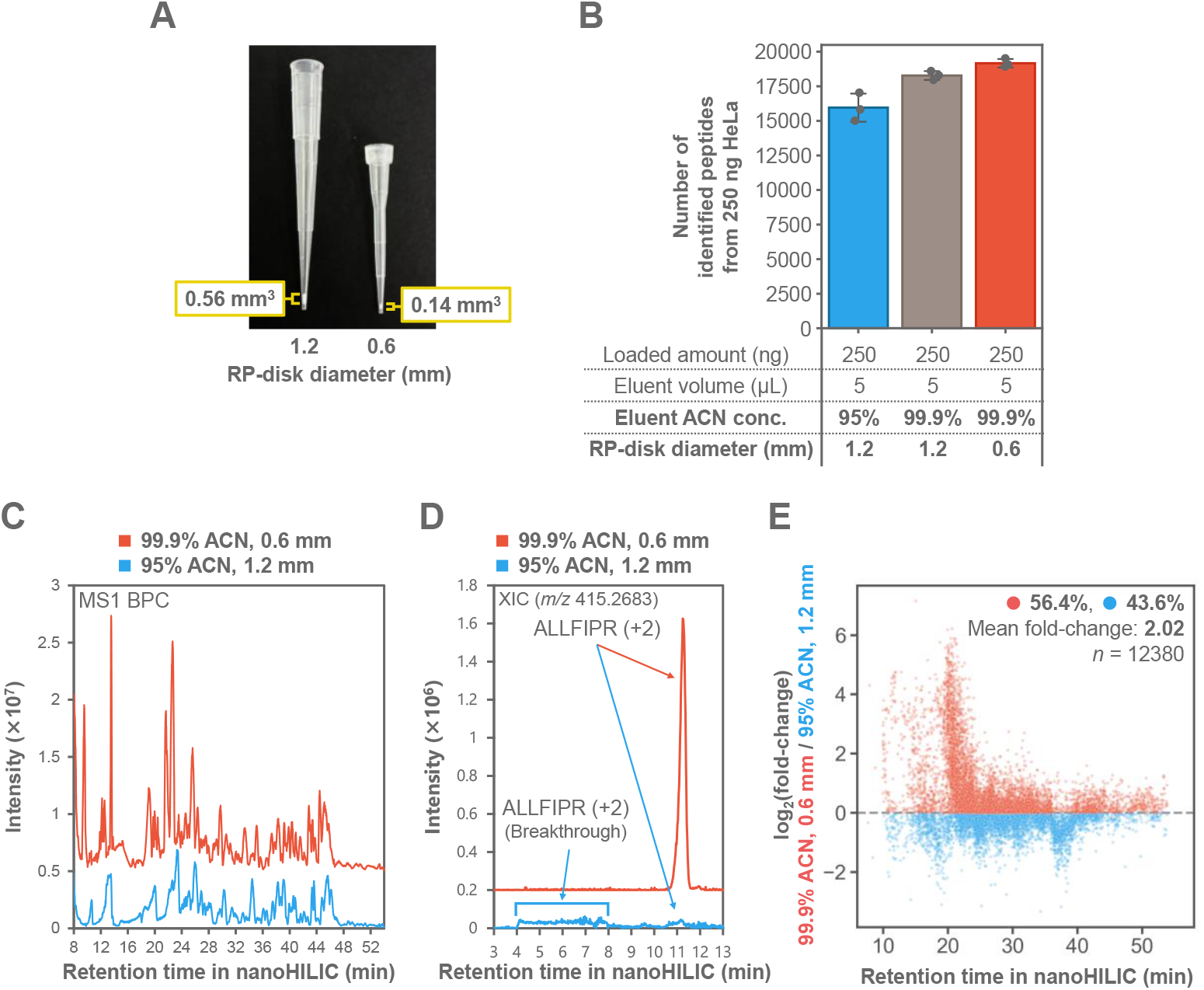
Optimization of DiReCT for entire-volume injection at the 5-µL scale. (A)Photographs of RP-StageTips packed with 1.2- or 0.6-mm RP disks and corresponding disk volumes. (B) Numbers of peptides identified under each condition. (C) MS1 base-peak chromatograms comparing the initial (1.2-mm disk, 95% ACN) and optimized (0.6-mm disk, 99.9% ACN) conditions. (D) Extracted ion chromatograms (m/z 415.2683, peptide ALLFIPR^2+^). (E) Log_2_(fold-change) in peptide intensities between optimized and initial conditions plotted against HILIC retention time. Red and blue dots indicate peptides with higher intensities under optimized and initial conditions, respectively. A total of 250 ng of HeLa peptides was desalted and eluted with 5 µL of 95% or 99.9% ACN containing 0.1% FA. The entire eluate was analyzed by nanoHILIC/MS/MS. Sample preparation was performed in triplicate.

Analysis of BSA digests confirmed that the initial 5 µL condition caused severe peak fronting and breakthrough, whereas the optimized condition restored peak widths and theoretical plate numbers to levels comparable to the 100 µL format (Figure S3).

### Benchmarking DiReCT/nanoHILIC/MS/MS Against nanoRPLC/MS/MS

To rigorously evaluate the analytical advantages of DiReCT/nanoHILIC/MS/MS, we benchmarked its performance against a matched nanoRPLC/MS/MS workflow across a wide range of peptide inputs (0.25–25 ng), corresponding approximately to the peptide content of 1–100 mammalian cells. This comparison was designed to isolate the chromatographic and ionization characteristics of each separation mode while ensuring that sample handling upstream of LC injection was optimized for the requirements of each method. Specifically, DiReCT enabled direct injection of the entire 5 µL RP-StageTip eluate under high-ACN conditions ideal for nanoHILIC, whereas nanoRPLC required drying and reconstitution in a low-organic solvent to ensure proper retention on the C18 column. Thus, the benchmarking experiment compared two workflows that were each optimized for their respective chromatographic modes, rather than imposing identical sample-solvent conditions that would disadvantage one method.

Across the entire dilution series, DiReCT/nanoHILIC/MS/MS demonstrated superior sensitivity at low input amounts (Figure 6). At peptide loads of 2.5 ng or less, nanoHILIC consistently identified more peptides and proteins than nanoRPLC, with the performance gap widening as the sample amount decreased. Most notably, from 0.25 ng of HeLa peptides—equivalent to the digest from approximately a single mammalian cell—DiReCT/nanoHILIC/MS/MS identified an average of 1177 peptides and 410 proteins. These values represent 8.9-fold and 6.7-fold increases, respectively, over the matched nanoRPLC/MS/MS workflow. This dramatic improvement highlights the suitability of nanoHILIC for trace-level proteomics when combined with a sample-introduction strategy that preserves peptide solubility and minimizes sample loss.

**Figure 6.**
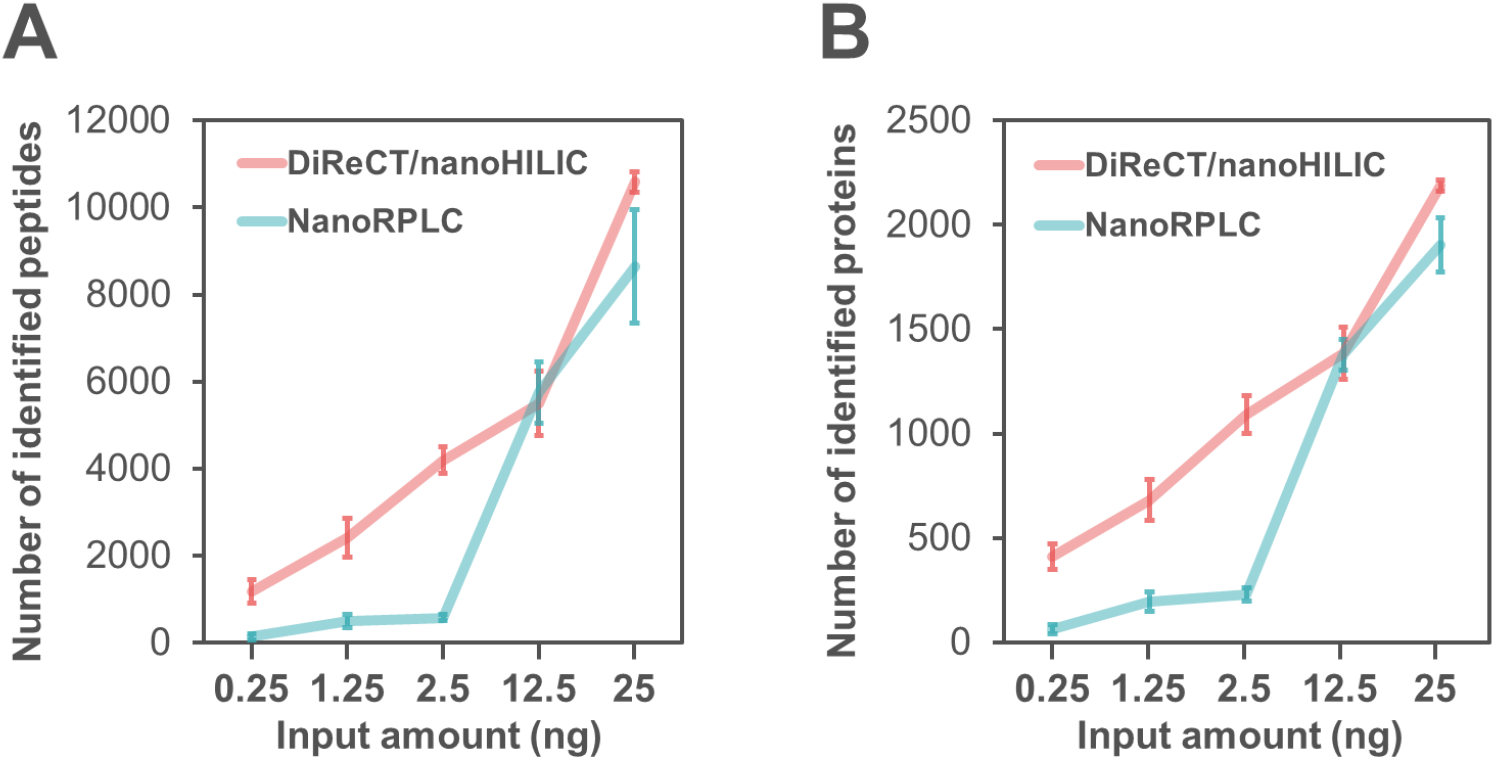
Benchmarking DiReCT/nanoHILIC/MS/MS against nanoRPLC/MS/MS for low-input proteomics. (A) Numbers of identified peptides. (B) Numbers of identified proteins. HeLa peptides (0.25–25 ng) were desalted using RP-StageTips packed with a 0.6-mm RP disk and eluted with 5 µL of 99.9% ACN containing 0.1% FA. For nanoHILIC/MS/MS, the entire eluate was injected directly. For nanoRPLC/MS/MS, eluates were dried and reconstituted in 5 µL of 4% ACN containing 0.5% TFA. NanoHILIC/MS/MS used a ZIC-HILIC column (3.5 µm, 100 Å, 200 mm × 100 µm i.d.), and nanoRPLC/MS/MS used a ReproSil-Pur 120 C18-AQ column (3 µm, 120 Å, 200 mm × 100 µm i.d.). Sample preparation was performed in triplicate.

Several factors likely contribute to the enhanced performance of DiReCT/nanoHILIC/MS/MS at low input levels. First, the ACN-rich solvent environment used throughout the DiReCT workflow reduces peptide adsorption to plastic surfaces, a well-known source of loss in low-abundance samples. Second, the high-organic mobile phase in nanoHILIC promotes efficient droplet desolvation and ionization in electrospray, improving MS sensitivity relative to aqueous-rich nanoRPLC conditions. Third, DiReCT eliminates the drying and reconstitution steps required in the nanoRPLC workflow, thereby avoiding losses associated with peptide precipitation or incomplete resolubilization. Finally, the orthogonal selectivity of HILIC may enhance the detectability of hydrophilic peptides that are poorly retained or suppressed in RPLC, further contributing to the increased identification depth.

Together, these results demonstrate that DiReCT/nanoHILIC/MS/MS provides a robust and highly sensitive platform for low-input proteomics. By integrating peptide solubilization, desalting, solvent exchange, and nanoHILIC-compatible elution into a single StageTip operation, DiReCT enables efficient sample transfer and maximizes the analytical advantages of nanoHILIC. The substantial gains observed at sub-nanogram peptide loads suggest that DiReCT/nanoHILIC/MS/MS is well suited for applications requiring high sensitivity, including single-cell proteomics, spatially resolved proteomics, and analyses of rare or precious biological samples.

## Conclusions

DiReCT provides a unified StageTip-based solution to the chained solvent-mismatch problem that has limited nanoHILIC/MS/MS for low-input proteomics. By coupling peptide solubilization, desalting, solvent exchange, and nanoHILIC-compatible elution within a single device, DiReCT enables efficient peptide recovery through a transient ACN gradient and eliminates drying-related losses. Conservative comparisons demonstrated that DiReCT matches or exceeds the performance of the two-step solubilization method, while optimization of StageTip geometry and eluent composition enabled entire-volume injection at the 5 µL scale with restored chromatographic performance. Under these optimized conditions, DiReCT/nanoHILIC/MS/MS substantially outperformed nanoRPLC/MS/MS at sub-nanogram peptide loads, establishing DiReCT as a practical and highly sensitive workflow for trace-level proteomics and a promising platform for single-cell and spatial proteomics applications.

## Author Contributions

Conceptual design of the project was done by E. Kanao and Y. Ishihama. The experimental designs were conducted by K. Akamatsu and E. Kanao. LC/MS/MS and data analysis were performed by K. Akamatsu. The manuscript was written by K. Akamatsu, E. Kanao, and Y. Ishihama.

## Acknowledgments

We would like to thank members of the Department of Molecular Systems BioAnalysis in Kyoto University for fruitful discussions. This work was supported by AMED-PRIME (JP25gm7010003h0001) to EK, AMED-Interstellar Initiative (JP25jm0610109h0001) to EK, JST-ASTEP (JPMJTR25U9) to EK, NEDO Intensive Support Program for Young Promising Researchers (seeds-4891) to EK, Grants-in-Aid for Scientific Research (KAKENHI; Grants JP26K02156 and JP23K13774 to EK, 25K22526, 23H04924, and 23K18185 to YI, 25KJ1530 to KA), Toyota Riken Scholarship to EK, and Takeda Science Foundation to EK.

## Associated Content

### Supporting Information

The Supporting Information is available free of charge at https://XXXX. It includes supplementary methods for the Q-Exactive acquisition used in Figure 3, gravimetric estimation of residual liquid in RP-StageTips and its effect on the ACN concentration of the final eluate, the effect of eluent ACN concentration on peptide identifications in the 100 µL-scale DiReCT workflow, and evaluation of nanoHILIC separation performance for BSA tryptic peptides prepared by DiReCT under different RP-StageTip conditions.

## Data availability statement

The MS raw data files and the analysis files containing the identification and quantification results for peptides and protein groups have been deposited at the ProteomeXchange Consortium (http://proteomecentral.proteomexchange.org) via the jPOST partner repository (https://jpostdb.org) with the dataset identifier JPST004538/PXD076566^.40,41^

---

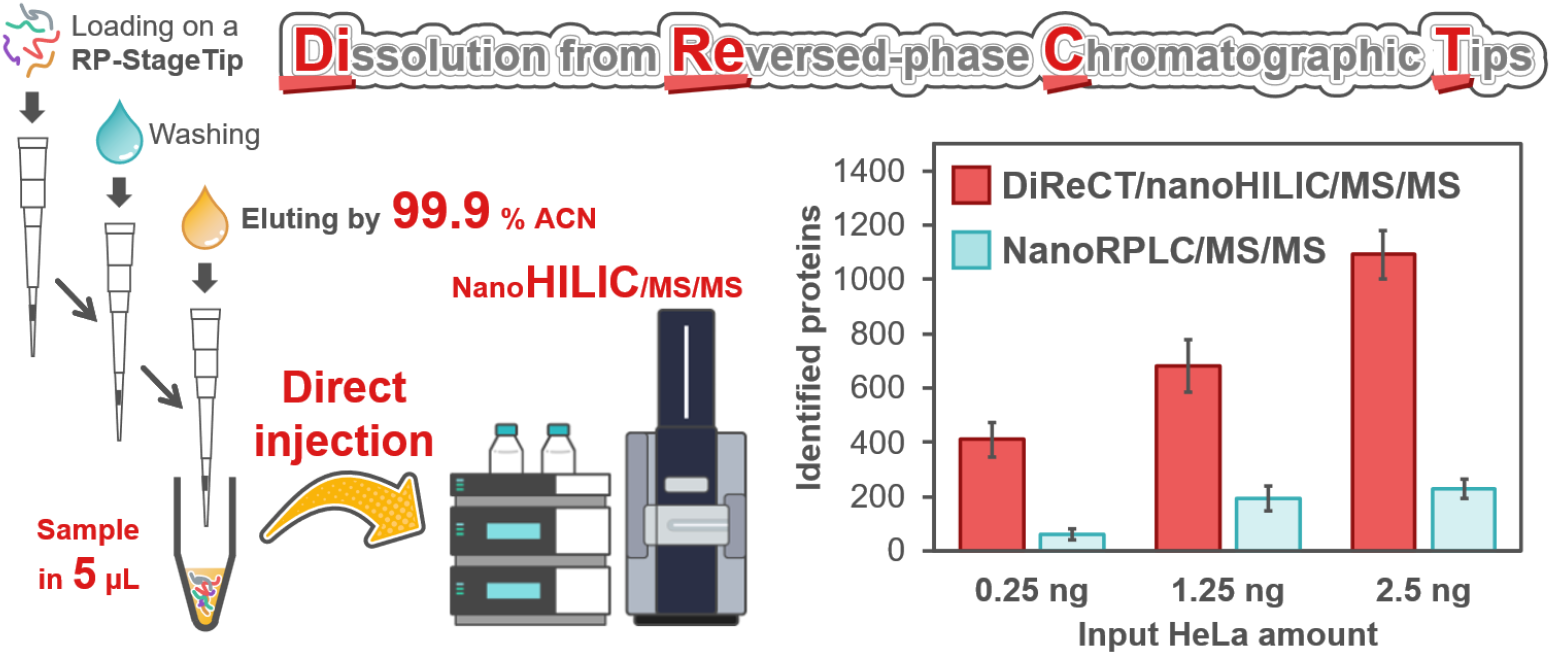

