## Supporting Information for "Direct Injection NanoHILIC/MS/MS Proteomics from Reversed-Phase StageTip Eluate"

1. **Supporting method**

2. **Supporting figures**

**Figure S1.** Gravimetric estimation of residual liquid in RP-StageTips and its effect on the ACN concentration of the final eluate.

**Figure S2.** Effect of eluent ACN concentration on peptide identifications in the 100-μL-scale DiReCT workflow.

**Figure S3.** Evaluation of nanoHILIC separation performance for BSA tryptic peptides prepared by DiReCT under different RP-StageTip conditions.

### 1. Supporting method

#### 1-1. Q-Exactive acquisition parameters

NanoHILIC/MS/MS analyses were performed on a Q-Exactive mass spectrometer (Thermo Fisher Scientific) coupled to an UltiMate 3000 pump and an HTC-PAL autosampler equipped with a 25 µL Hamilton X-Type syringe. Samples were maintained at 4 °C in the autosampler. Peptides were separated on a self-pulled needle column (200 mm × 100 µm i.d.) packed with ZIC-HILIC particles (3.5 µm, 100 Å, Merck) at a flow rate of 1000 nL/min and a column temperature of 20 °C. Mobile phase A was 0.1% FA in ACN, and mobile phase B was 0.1% FA in water. The gradient was: 0–5 min, 5–20% B; 5–25 min, 20–35% B; 25–40 min, 35–60% B; 40–45 min, 60–70% B; 45–45.1 min, 70–5% B; and 45.1–60 min, 5% B. The injection volume was 5 µL.

The electrospray voltage was set to 2.4 kV in positive mode. Survey scans were acquired from m/z 350–1500 at a resolution of 70,000 with an AGC target of $3 \times 10^6$ and a maximum injection time of 100 ms. MS/MS spectra were acquired using a Top10 method at a resolution of 15,000 with an AGC target of $1 \times 10^5$ and a maximum injection time of 100 ms. Fragmentation was performed by HCD with a normalized collision energy of 27%, and dynamic exclusion was set to 30 s.

Tryptic peptides from HeLa lysate were analyzed using FragPipe (v23.1) with MSFragger (v4.3) and IonQuant (v1.11.11) against the UniProtKB/Swiss-Prot human database (November 2024, 20,429 entries). Precursor and fragment mass tolerances were set to 20 ppm. Trypsin/P was specified as the enzyme with up to two missed cleavages. Carbamidomethylation of cysteine was set as a fixed modification, and methionine oxidation and protein N-terminal acetylation were set as variable modifications. MSBooster rescoring used spectral and ion-mobility predictions (retention-time prediction disabled). Results were filtered to a 1% FDR at both the PSM and protein levels.

#### 2. Supporting figures

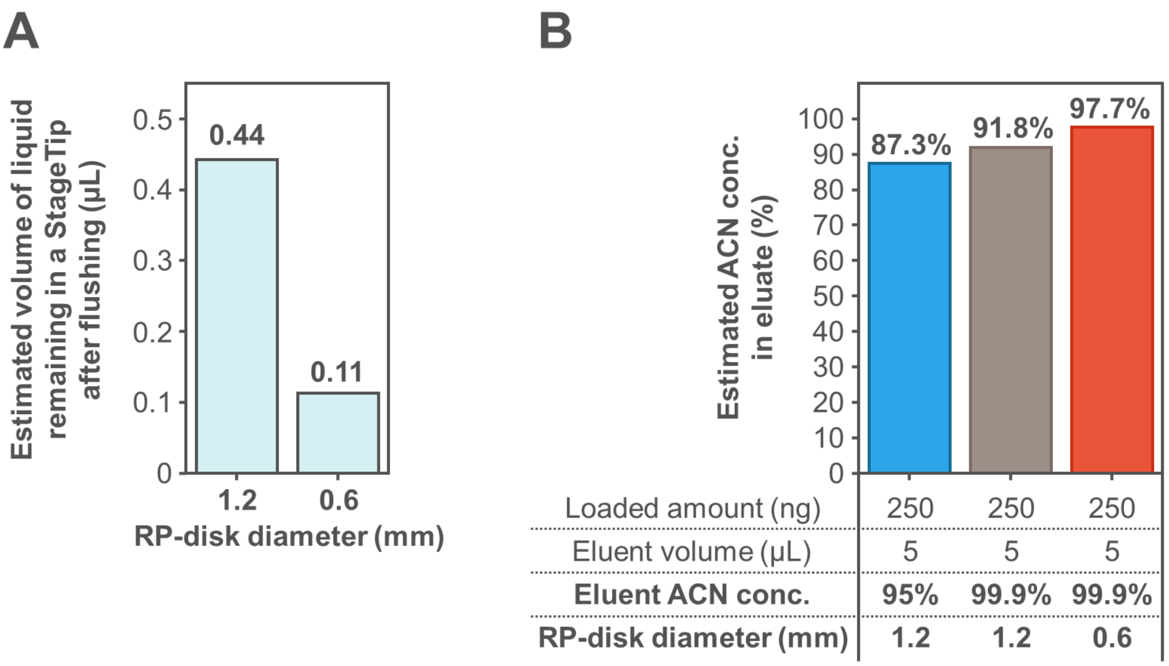


**Figure S1. Gravimetric estimation of residual liquid in RP-StageTips and its effect on the ACN concentration of the final eluate.**

(A) Estimated volume of liquid remaining in RP-StageTips after washing for each RP-disk size. (B) Estimated ACN concentration in the final eluate after mixing of the residual liquid with the eluent under each condition. The estimated ACN concentration in the eluate was calculated as follows: Estimated ACN concentration in eluate (%) = [Eluent ACN concentration (%) × Eluent volume (µL)] / [Eluent volume (µL) + estimated volume of liquid remaining in a StageTip after flushing (µL)].

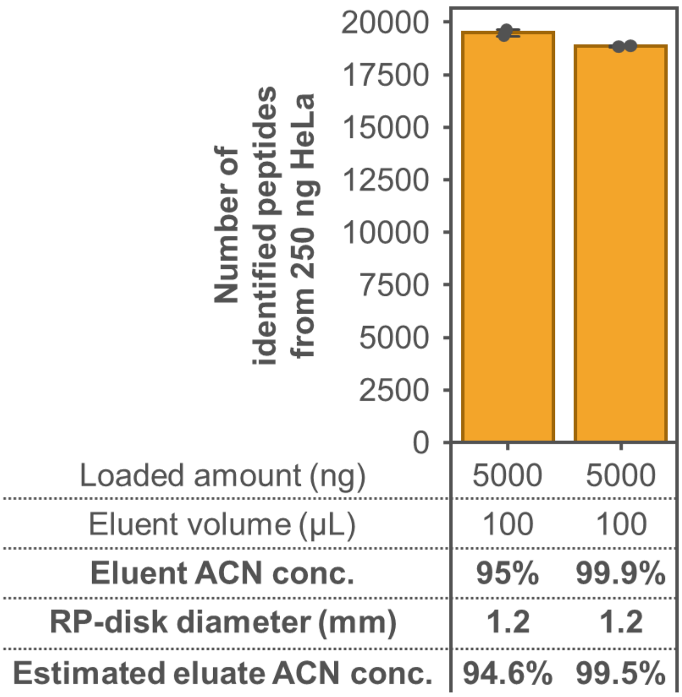


**Figure S2. Effect of eluent ACN concentration on peptide identifications in the 100-μL-scale DiReCT workflow.**

A total of 5000 ng of HeLa peptides was desalted using an RP-StageTip packed with a 1.2 mm RP-disk punched with a 16-gauge needle and then eluted with 100 μL of solvent containing either 95% or 99.9% ACN and 0.1% FA. A 5-μL aliquot of the eluate, corresponding to 250 ng of peptides, was analyzed by nanoHILIC/MS/MS. Sample preparation was performed in duplicate. The estimated ACN concentration in the final eluate was calculated using the same equation as in Figure S1.

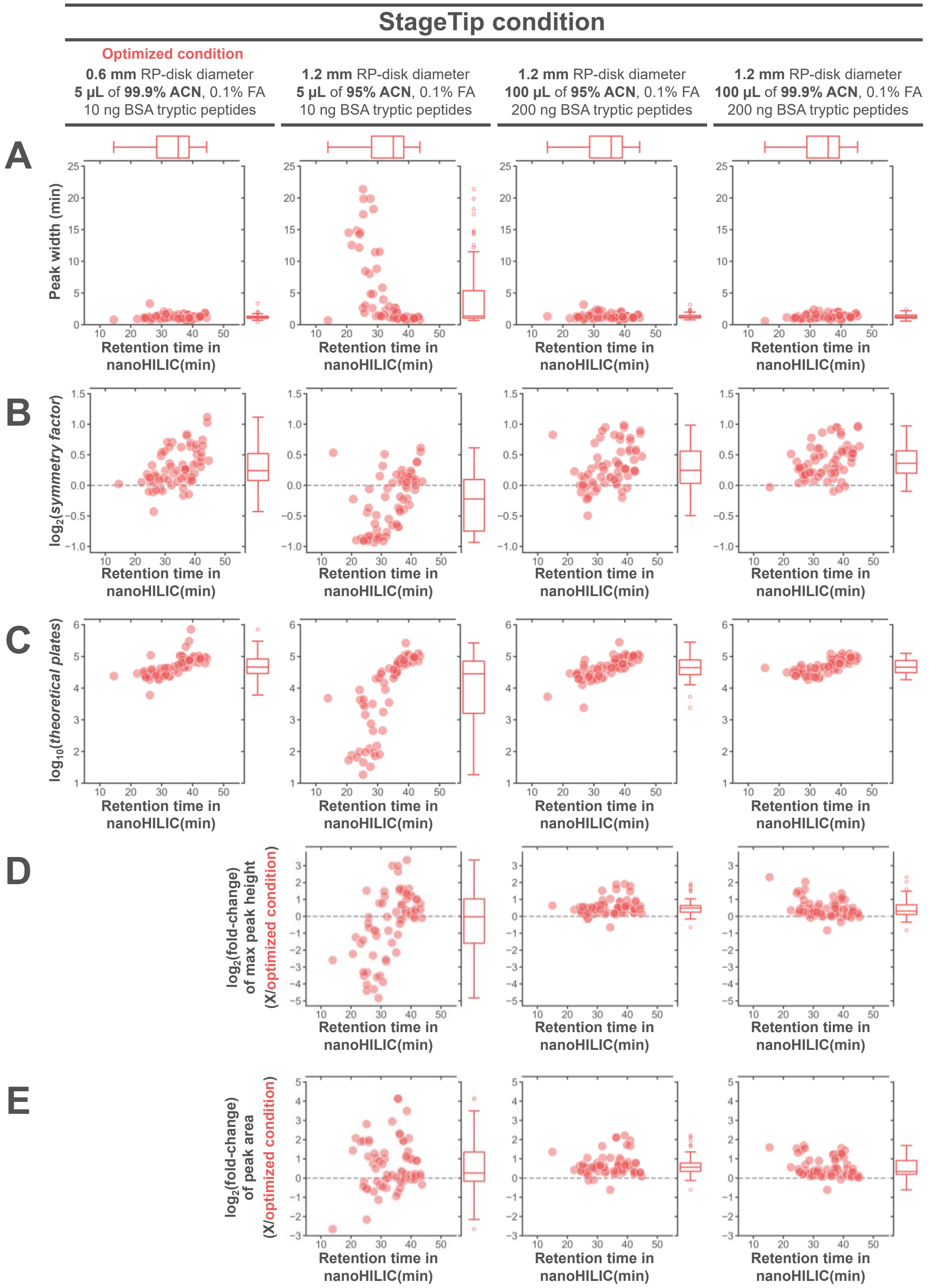


**Figure S3. Evaluation of nanoHILIC separation performance for BSA tryptic peptides prepared by DiReCT under different RP-StageTip conditions.**

BSA tryptic peptides (10 or 200 ng) were desalted using RP-StageTips packed with RP-disks punched with either a 16- or 20-gauge needle and then eluted with 5 or 100 μL of solvent containing either 95% or 99.9% ACN and 0.1% FA. For the 5 μL-scale conditions, the entire eluate was analyzed directly by nanoHILIC/MS/MS. For the 100-µL-scale conditions, a 5-µL aliquot of the eluate, corresponding to 10 ng of peptides, was analyzed. (A–E) Plots of peptide-ion properties against HILIC retention time: (A) peak width, (B) $\log_2$(symmetry factor), (C) $\log_{10}$(theoretical plate number), (D) $\log_2$(fold change in maximum peak height relative to the optimized condition), and (E) $\log_2$(fold change in peak area relative to the optimized condition).
